# The critical modulatory role of spiny stellate cells in seizure onset based on dynamic analysis of a neural mass model

**DOI:** 10.1101/2021.07.19.452876

**Authors:** Saba Tabatabaee, Fariba Bahrami, Mahyar Janahmadi

## Abstract

Growing evidence suggests that excitatory neurons in the brain play a significant role in seizure generation. Nonetheless, spiny stellate cells are cortical excitatory non-pyramidal neurons in the brain, which their basic role in seizure occurrence is not well understood. In the present research, we study the critical role of spiny stellate cells or the excitatory interneurons (EI), for the first time, in epileptic seizure generation using an extended neural mass model inspired by Liu and Wang model in 2017. Applying bifurcation analysis on this modified model, we investigated the rich dynamics corresponding to the epileptic seizure onset and transition between interictal and ictal states caused by EI connectivity to other cell types. Our results indicate that the transition between interictal and ictal states (preictal signal) corresponds to a supercritical Hopf bifurcation, and thus, the extended model suggests that before seizure onset, the amplitude and frequency of neural activities gradually increase. Moreover, we showed that 1) the altered function of GABAergic and glutamatergic receptors of EI can cause seizure, and 2) the pathway between the thalamic relay nucleus and EI facilitates the transition from interictal to the ictal activity by decreasing the preictal period. Thereafter, we considered both sensory and cortical periodic inputs to study model responses to various harmonic stimulations. Bifurcation analysis of the model, in this case, suggests that the initial state of the model might be the main cause for the transition between interictal and ictal states as the stimulus frequency changes. The extended thalamocortical model shows also that the amplitude jump phenomenon and nonlinear resonance behavior result from the preictal state of the modified model. These results can be considered as a step forward to a deeper understanding of the mechanisms underlying the transition from normal activities to epileptic activities.

## 1. Introduction

Epilepsy is one of the most common disorders of the Central Nervous System (CNS), which causes sudden abnormal and synchronized brain activities, resulting in seizures. Interactions between excitatory and inhibitory neurons shape mainly brain activities and some transitory disparity in the inhibitory/excitatory balance can trigger a seizure. One-third of people with epilepsy are likely to have drug resistant epilepsy, and treatments such as surgery or stimulation-based treatments like deep brain stimulation (DBS) can be considered for these kinds of drug-resistant patients [1]. A seizure can be composed of four distinct states including preictal, ictal, interictal and postictal. The preictal state appears before the seizure begins and indicates that seizures do not simply start out of nothing. Experimental studies [2] and [3] on human tissue and intracranial EEG signals have shown that preictal spikes are distinguishable from ictal and interictal signals. The preictal signal can be used to predict seizure occurrence. Understanding the mechanism of generation of the preictal period opens the way for a more precise seizure prediction and thereby more reliable automatic interventions to prevent the seizure occurrence [4].

Different seizures can be categorized into focal and generalized onset seizures. Tonic, clonic and absence seizures are the generalized onset types of seizures and they occur when a widespread activity triggers in both hemispheres of the brain [5]. Furthermore, there are two groups of seizures that have known hemispheric origins, *i.e.,* generalized and focal onset seizures [6]. Both groups are categorized into two sub-classes: motor and non-motor seizures. Both sub-classes have several motor seizure types in common (e.g., tonic and clonic seizures). However, each type has different manifestations and symptoms. On the other hand, absence seizures are among well-known non-motor generalized onset seizure groups that are also considered as sensory and emotional seizures [6]. In tonic-clonic seizures, a clonic activity follows the tonic activity. Electrophysiological observations have shown the two-way transition between absence and tonic-clonic seizures [7] and dysfunction of the cortical and/or thalamic circuitries is believed to produce the absence, clonic and tonic epileptic activities, together with transitions between them. Recorded electroencephalogram (EEG) signals indicate that seizures can vary in frequency contents. A typical absence seizure can be considered as approximately synchronous spike and wave discharges (SWD) with a frequency of 2~4 Hz, but, atypical absence seizures (another kind of non-motor generalized onset seizures) show different frequency ranges (< 2 Hz and > 4 Hz) [8]. A tonic seizure is a fast-spiking activity (> 14 Hz) with low amplitude, and a clonic seizure is slow-wave activity (approximately 3 Hz) with high amplitude.

Several studies have shown that interneurons are cell types, which play key roles in the initiation and termination of epileptic seizures [9, 10]. Interneurons in the cortex, not only control the activity of pyramidal neurons, but they also receive thalamic relay nucleus inputs. Therefore, they are important factors in transferring and integrating the sensory information coming from the thalamus to the cortex [11]. Interneurons are traditionally regarded as inhibitory neurons; but more precisely, there are two kinds of interneurons in the CNS; *i.e.*, excitatory and inhibitory interneurons. Inhibitory interneurons use the neurotransmitter gamma-aminobutyric acid (GABA) or glycine, and excitatory interneurons (EI) are spiny stellate cells in the neocortex of the human brain and use glutamate as their neurotransmitters [12]. A study on the human brain [13] has shown that the neurons in the neocortex play a crucial role in SWD. Another study on the juvenile mice using multi-photon imaging [14] suggested that the neocortex has an intrinsic predisposition for seizure generation and pathological recruitment of the thalamus into joint synchronous epileptic activities. Genetic mutation study on mice has shown that in the spiny stellate cells (in the neocortex) an alteration in the kinetic of N-methyl-D-aspartate receptors (NMDA) was sufficient to cause neuronal hyper-excitability leading to epileptic activity in the brain [15]. An experimental study on monkeys has shown that synchronous discharges of excitatory interneurons could spread the epileptic activity in the brain, and EI could play an important role in the initiation and propagation of SWD, in which the spike is because of the extracellularly synchronous and powerful depolarization of excitatory interneurons, and the wave is because of the inhibitory interneurons. Therefore, SWD can be generated as a result of the interactions between inhibitory and excitatory interneurons [16]. Nevertheless, it is not well understood how excitatory interneurons in the neocortex of the human brain can explicitly cause a seizure. Therefore, understanding the epileptogenic role of the excitatory interneurons in the neocortex of the human brain is crucial to deepen our knowledge about neocortical epilepsy.

On the other hand, computational modelling of epilepsy has provided dynamic insights into the mechanism underlying the transition from normal to epileptic activities. Fan et al. [17] used a modified computational field model of a cortical microzone and showed that the two-way transitions between absence and tonic-clonic epileptic seizures are induced by disinhibition between slow and fast inhibitory interneurons. Nevertheless, they ignored the role of thalamic circuitry and its interaction with the cerebral cortex in the proposed computational model; whereas, in [18, 19] it is shown that the mechanism of seizures depends on the dysfunctionality in the function of the thalamus and or cortex. Experimental evidence suggests that interactions between the thalamus and cerebral cortex influence the initiation and propagation of SWD [20]. For that reason, Taylor et al. [21] developed a thalamocortical model to investigate the SWD generation. Liu and Wang [22] introduced a thalamocortical model inspired by the model developed by Taylor and colleagues [21] and Fan et al. [17]. In their mode,l they considered the dual pathways between fast and slow inhibitory interneurons in the cortex and simulated normal activity, clonic, absence and tonic seizures. However, none of the models mentioned above did consider the excitatory interneurons population in the thalamocortical circuitry, and therefore, they did not study the role of EI population in transition from the interictal to the ictal state.

In the present study, we modified and extended a thalamocortical model, originally proposed by Liu and Wang [22]. The original model describes absence, tonic, clonic and tonic-clonic seizures generation. That is one of the reasons we chose this model and extended it to investigate the role of excitatory interneurons on seizure onset using dynamical analysis. For this purpose, we introduced an additional group of interneurons into the model, which was excitatory and considered their interactions with pyramidal neurons, inhibitory interneurons and thalamic relay nucleus in the brain. As will be shown, this model is not only capable of producing different types of epileptic seizures such as absence, clonic and tonic, but, it also generates the preictal period.. The original model of Liu and Wang [22] does not describe the preictal period.

The organization of the paper is as follows. in section 2, the modified epileptic dynamical model inspired by the model proposed in [22] is introduced. Then, in section 3, we first explore the role of impairment in the activity of the excitatory interneurons in intericatl to ictal transition. Using the modified model, we investigate the role of GABAergic and glutamatergic receptors of excitatory interneurons in seizure generation and transition between interictal and ictal states. Through numerical simulations and bifurcation analysis, we show that dysfunction of excitatory and inhibitory receptors of excitatory interneurons leads to interictal to ictal transition. Moreover, we investigate the effect of impairments in the interactions between the thalamic relay nucleus and excitatory interneurons and we will investigate how they give rise to interictal to preictal transition, and also facilitate the occurrence of an epileptic seizure. Then, we investigate the behavior of the extended model in a more general framework when the excitatory interneurons function properly, but there are dysfunctions in the thalamus, more exactly, in the synaptic strength between the thalamic relay nucleus and reticular nucleus. Finally, in section 4, we examine the frequency responses of the modified thalamocortical model subjected to various sensory and cortical periodic inputs to identify the effect of a various stimuli on epileptic seizures. The obtained results show that the model starts its nonlinear resonator behavior when the shift from normal activity to preictal state takes place. Therefore, the amplitude jump phenomenon corresponding to chaotic behavior takes place before the onset of a seizure.

## 2. Materials and Methods

### 2.1. Model Structure

Recent studies [23,24] have shown that during generalized seizures, dysfunctionality has been identified in the thalamocortical network. Therefore, both the cortex and thalamus play an important role in seizure generation. Accordingly, we modified the thalamocortical model of Liu and Wang [22] and as shown in Fig. 1. This model is a neural mass model, which considers the inhibitory and excitatory neuronal populations and their connections in the brain. The cortical section of this model includes excitatory pyramidal neuron population (PY), mutual fast and slow inhibitory interneuron populations (I1, I2) having inhibitory GABA_A_ and GABA_B_ receptors. Activation of ionotropic GABA_A_ receptors causes fast inhibitory postsynaptic potentials (IPSPs) by allowing the influx of Cl^-^ into the postsynaptic cells, while activation of metabotropic GABA_B_ receptors mediates a slow inhibition by inducing K^+^ efflux The thalamic subsystem includes a population of excitatory thalamic relay nucleus (TC) and the inhibitory population of neurons located in the reticular nucleus (RE). To investigate the effect of spiny stellate cells on seizure onset, we considered the excitatory interneurons population, EI, in our thalamocortical model (see Fig 1). In this new model version, we will investigate the synaptic connectivity strength of excitatory interneurons to explore dynamics, which lead to preictal state and dynamics during transitions between absence, tonic and clonic seizures.

**Fig. 1.**
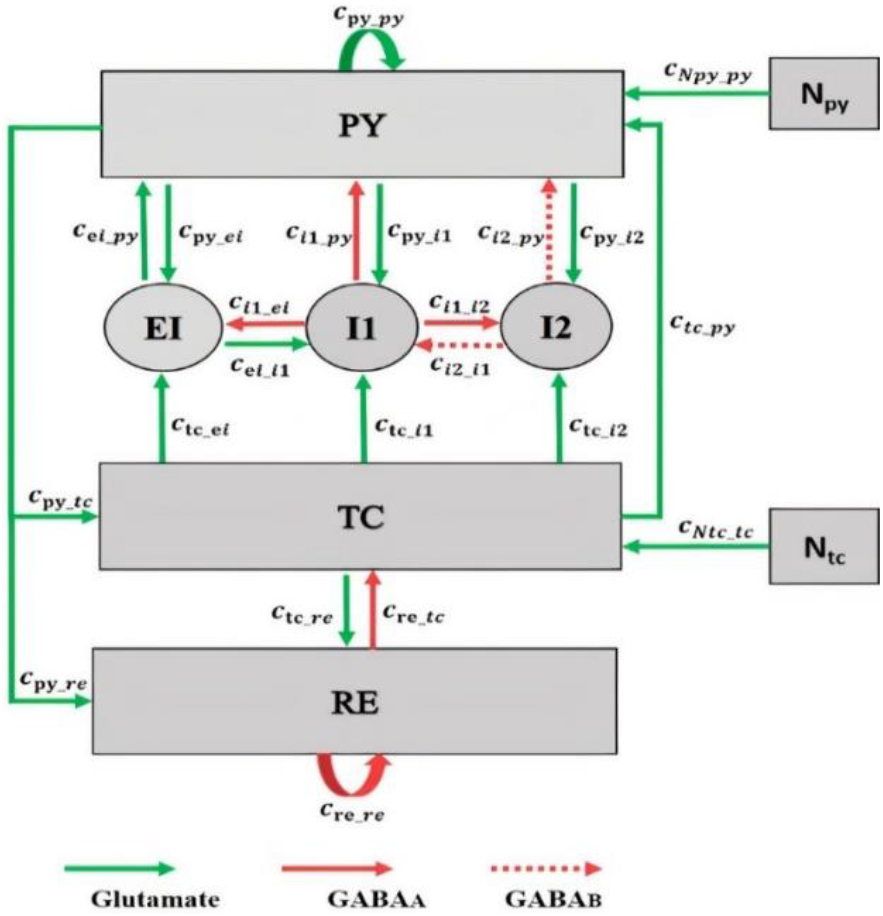
Scheme of the thalamocortical model including the excitatory interneuron population in the cortex. The model involves cortical and thalamic subnetworks. In the cortical subnetwork, PY is the pyramidal neuronal population, EI is the excitatory interneurons population, I1 and I2 are the fast and the slow inhibitory interneurons populations, respectively. In the thalamic subnetwork, TC is the thalamic relay nucleus population and RE is the thalamic reticular nucleus population. Green arrows define the excitatory glutamatergic receptor. Red solid and dashed arrows define the GAGA_A_ and GABA_B_ inhibitory receptors, respectively. The value of parameters of this model are: c_py_py_ = 1.89, c_py_i1_ = 4, c_i1_py_ = 1.8, c_re_re_ = 0.01, c_tc_re_ = 10, c_re_tc_ = 1.4, c_py_tc_ = 3, c_py_re_ = 1.4, c_tc_py_ = 1, c_py_i2_ = 1.5, c_tc_i1_ = 0.05, c_tc_i2_ = 0.05, c_ei_i1_ = 0.05, c_ei_py_ = 0.442, c_i2_py_ = 0.05, c_i2_i1_ = 0.1, c_i1_i2_ = 0.5, c_Npy_py_ = 1, c_Ntc_tc_ = 1, tau_1_ = 21.5, tau_2_ = 31.5, tau_3_ = 0.1, tau_4_ = 4.5, tau_5_ = 3.8, tau_6_ = 3.9, h_py_ = −0.4, h_i1_ = −3.4, h_i2_ = −2, h_ei_ = −1, h_tc_ = −2.5, h_re_ = −3.2, ε = 250000, a_tc_=0.02, a_py_=0.02, B_Npy_=0.7 and B_Ntc_=0.1. These parameters were obtained by trials and error such that the dynamic behavior of the model (*i.e.,* the bifurcation diagrams) were the same as the original model developed by Liu and Wang [22].

In this thalamocortical model, we analyze the interactions between neuronal populations in the network by varying the strength of the synaptic connections. These synaptic connections are associated with the type of receptors. In the brain, receptors are either excitatory (glutamatergic) or inhibitory (GABAergic). In the cortex, the pyramidal population is excitatory n and there is a mutual excitatory connection between pyramidal and excitatory interneurons by the glutamatergic receptors [25]. Excitatory and inhibitory interneurons are locally [26] and based on this fact, we consider an excitatory and an inhibitory connection between excitatory interneurons and fast inhibitory interneurons. Thalamus is globally connected to the excitatory interneurons by the thalamic relay nucleus, and the thalamic relay nucleus is regarded as an excitatory population [27]. Here, we consider an excitatory connection from the thalamic relay nucleus population to the excitatory interneurons.

The thalamic relay nucleus receives all the sensory information from different parts of the brain and then this information is sent to the appropriate area in the cortex for further processing [28,29]. Here, we investigate the dynamic of the extended model when it receives cortical and sensory inputs. In this regard, we consider an input (N_tc_) to the thalamic relay nucleus by adding a periodic sensory input with frequency ftc, bias B_Ntc_ and amplitude a_tc_. Moreover, we consider a cortical input (N_py_) to the pyramidal neuronal population by adding a biased sinusoidal waveform with bias of B_Npy_, frequency of fpy and amplitude of a_py_.

We implement the model using equations developed by Liu and Wang [22] and the Amari neural field equations [30] (for the EI population). The differential equations of the model are given below.

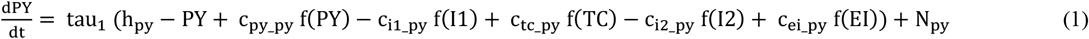

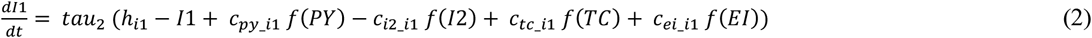

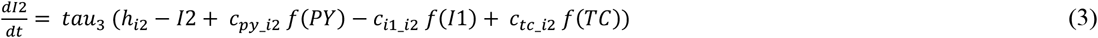

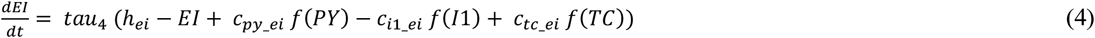

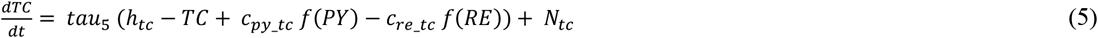

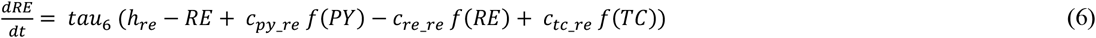

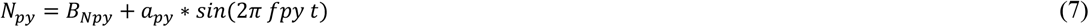

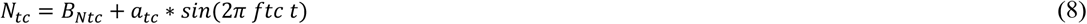

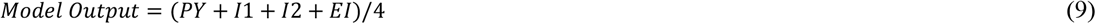

Dimensionless parameters c_1,…,16,iny,ei,in1,in2_ are the connectivity parameters, which determine the coupling strength between the populations, and h_py,i1,i2,ei,tc,re_ are input parameters, tau_1,2,…,6_ are time scales parameters, f(x) = ^1^/(1+ε^-x^) is the transfer function, where ε determines steepness and x=PY,I1,I2,EI,TC and RE. N_py_ (cortical input) and N_tc_ (sensory input) are inputs to the pyramidal population and thalamic relay nucleus population, respectively. These parameters were obtained by trials and error such that the dynamic behavior of the model (*i.e*., the bifurcation diagrams) has similar mechanism as model developed by Liu and Wang [22]. The output of the model is taken as the mean activity of the four cortical populations.

### 2.2. Simulations

Simulations are performed by the standard fourth-order Runge-Kutta integration using MATLAB 9.4 and the MatCont environments, with a step size of 0.0039 s. We set the time window at 60 s, and use the last two seconds of the time series to analyze the stable state of the time domain and deterministic behaviour of the model. We use the frequency domain and time domain analysis to explore the transitions from interictal to ictal state. We extract dominant frequency (the frequency that carries the maximum energy) from the power spectral density using the fast Fourier transform, and for the bifurcation analysis, we extract extrema (local maximum and minimum) of the mean of four cortical populations from the time series. By doing so, we can observe seven different epileptic activities such as interictal state (or the normal background activity), preictal state, which occurs before the seizure onset, slow rhythmic activity that can be observed at seizure onset or during the seizures, typical absence seizures, atypical absence seizures, tonic and clonic seizures. For 1D bifurcation diagrams (section 3.1.1, 3.1.2 and 3.1.4) we obtain the maximum and minimum of model output as we change the bifurcation parameter. For hybrid bifurcation analysis (section 3.1.3) when both of the parameters of c_py_ei_ and c_i1_ei_ are changed, we computed the local maxima, local minima and frequency of the last two seconds of the model output, and then, we sorted the local maxima and minima from large to small to obtain the largest (Pmax1) and the smallest (Pmax2) maxima, the largest (Pmin1) and the smallest (Pmin2) minima. We summarized our categorization in Table 1.

**Table 1.** Specific characteristics of seven simulated signals. Dominant frequency, the largest minima, the largest maxima, the largest minima and the largest maxima are characteristics used to separate seven different mean activities of the cortical populations.

| Type of the signal | Dominant frequency range | The largest (Pmin1) and the smallest minima (Pmin2) | The largest (Pmax1) and the smallest maxima (Pmax2) |
| --- | --- | --- | --- |
| Normal background activity | DF = 0 Hz | - | - |
| Preictal spikes | DF < 3.5 Hz | Pmin1-Pmin2<0.2 | 0.01<Pmax1-Pmax2<0.12 |
| Slow rhythmic activity | DF < 15 Hz | - | -0.8<Pmax1 <-0.1 |
| Typical absence seizure | DF ∈ [2,4] Hz | 0.004<Pmin1-Pmin2 | - |
| Atypical absence seizure | DF > 4 Hz and DF < 2 Hz | 0.01 <Pmin1-Pmin2 | - |
| Clonic seizure | DF ∈ [2,4] Hz | Pmin1-Pmin2<0.15 | 0.01<Pmax1-Pmax2 |
| Tonic seizure | DF ≥ 14 Hz | Pmin1-Pmin2<0.01 | 0.01<Pmax1-Pmax2 |

## 3. Results

In this section, the role of excitatory interneurons in transition from interictal to ictal, and the frequency response of the thalamocortical model, when it receives sensory and cortical inputs, are examined using various bifurcation analysis and dynamical simulations. In section 3.1, to explore the transition dynamics of the model we assume that the inputs of model are constant; *i. e.*, N_py_ = B_Npy_ and N_tc_ = B_Ntc_. In section 3.2, for investigating the frequency response of the model, a sinusoidal waveform signal is added to both sensory and cortical inputs, that is, N_py_ = B_Npy_ + a_py_ * sin(2π fpy t) and N_tc_ = B_Ntc_ + a_tc_ * sin(2π ftc t).

### 3.1. Transition dynamics

Excitatory interneurons are connected to the pyramidal neurons and fast inhibitory interneurons by glutamatergic and GABAergic receptors, respectively. The synaptic strength is not static and the changes in the neurotransmitters released by the excitatory and inhibitory neurons in the brain can result in short term or long term changes in synaptic strength. Antiepileptic drugs either block glutamatergic receptors or facilitate the function of GABAergic receptors, which result in a change in the glutamatergic and GABAergic neurotransmitter release [31,32]. Moreover, a ketogenic diet can bring about the altered function of receptors and change in neurotransmitter release [33]. Experimental evidences have shown that, changes in the function of GABAergic and glutamatergic receptors in the cortex of rats can be seen in genetic absence, tonic and clonic seizures onset [34,35,36]. However, the transition dynamics from interictal to ictal, and transitions between absence, tonic and clonic seizures caused by the abnormalities in glutamatergic and GABAergic receptors of EI are still unclear. In this section, by using bifurcation analysis, we explore the effect of impairment in the function of GABAergic and glutamatergic receptors of EI population on the epileptic dynamics of the model as the synaptic strengths c_i1-ei_ and c_py-ei_ change, respectively.

#### 3.1.1. Transition dynamics produced by GABAergic receptor-mediated inhibition in EI

Increasing the GABAergic inhibition is traditionally believed to suppress epileptic seizures, however, a computational study has shown that before seizure onset, the activity of GABAergic interneurons increases that leading to synchronous neuronal activity and epileptic seizures [37]. Based on an experimental study on human, antiepileptic drugs such as Midazolam that increase the GABAergic neurotransmitters in the brain can trigger seizure [38]. Abnormality in GABA_A_ receptors brings about a shift in the chloride reversal potential in neurons, which in turn results in changing the behavior of GABA from inhibitory to excitatory behavior and cause seizure [39]. Therefore, the role of GABAergic receptors epileptic seizures is more complex than one could assume that they have only inhibitory roles and anticonvulsant agents always inhibit their function. Here, we investigate the effect of increasing inhibitory function of GABAergic receptors of excitatory interneurons in the neocortex on seizure generation. We explore the epileptiform activity induced by the dysfunction of the GABAergic receptor of the excitatory interneuron population as the synaptic strength c_i1-ei_ changes. Results of Fig. 2 show that the model displays rich dynamics as the parameter c_i1-ei_ varies.

**Fig. 2.**
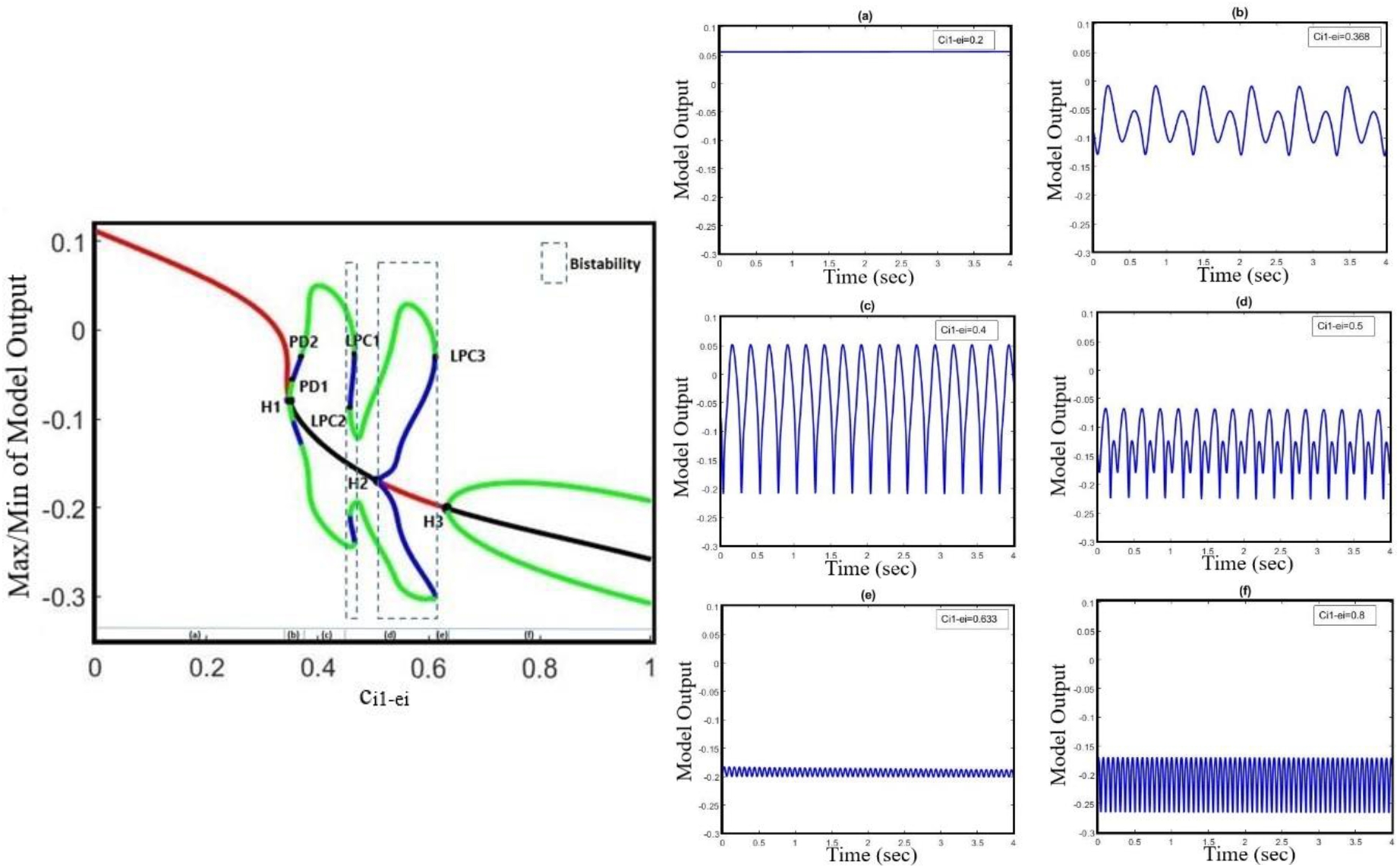
Bifurcation diagram and corresponding time series of the model output for different values of c_i1_ei_. The model output is defined as the mean value of output voltage of PY, I1, I2 and EI populations. The bifurcation diagram of the model (left) is calculated and plotted for c_i1_ei_ as the bifurcation parameter and with c_py_ei_ =0.8, c_tc_ei_ =4.5, a_tc_=0 and a_py_=0. we also set the a_tc_=0 and a_py_=0. In the plot, blue and green cycles, represent the unstable and stable limit cycles, respectively, and red and black lines, represent the stable and unstable equilibrium points, respectively. According to the diagram as the parameter ci1_ei changes, the model produces (a) normal background firing, (b) pre-ictal spikes, (c) clonic discharges, (d) SWD, (e) slow rhythmic activity and (f) tonic discharges. LPC1, LPC2 and LPC3 are fold limit cycle bifurcation, H1 and H3 are supercritical Hopf bifurcation, H2 is subcritical Hopf, and PD1 and PD2 are period of doubling bifurcations.

In Fig. 2, the overall bifurcation is shown in detail for variations of c_i1-ei_ as the bifurcation parameter. In the bifurcation diagram from left to right, the red line corresponds to the stable fixed points, which are related to the normal background activity (Fig. 2(a)), and upon increasing the c_i1-ei_ to ~ 0.349 a stable fixed point coalesces with a stable limit cycle and a supercritical Hopf bifurcation point (H_1_) happens, with increasing the c_i1-ei_ to ~ 0.355 a period doubling bifurcation (PD1) occurs, which forms the preictal spikes patterns (transition from normal background activity to ictal activity, Fig. 2(b)). With a further increase of c_i1-ei_ parameter to ~ 0.37, another period doubling bifurcation (PD2) happens and the transition from preictal state to clonic seizure takes place and shapes clonic seizure patterns (Fig. 2(c)). In addition, as c_i1-ei_ is increased up to ~ 0.458, the first fold limit cycle bifurcation (LPC1) occurs, and another fold limit cycle bifurcation (LPC2) takes place with further increase in c_i1-ei_. With the coexistence of two stable limit cycles in the range of c_i1-ei_ bounded by two-fold limit cycles bifurcations (LPC1 and LPC2) a bistable region appears and creates the coexistence of two different amplitude SWD patterns (Fig. 2(d)). Moreover, stable points appear again at c_i1-ei_ around 0.508, where the first subcritical Hopf bifurcation (H_2_) takes place, and another bistable region turns out with coexistence of stable fixed points and a stable limit cycle until the third fold limit cycle bifurcation (LPC3) at c_i1-ei_ ~ 0.611 takes place. Upon increasing the c_i1-ei_, the stable focus points, which are corresponding to slow rhythmic activity take place (Fig. 2(e)), and then at c_i1-ei_ ~ 0.634 another supercritical Hopf bifurcation (H_3_) happens and shapes the tonic seizure patterns (Fig. 2(f)).

#### 3.1.2. Transition dynamics produced by glutamatergic receptor-mediated excitation in EI

Studies have shown that antiepileptic drugs that are expected to reduce excitation in the brain, on the contrary, can have a paradoxical effect and brings about an aberrant synchronization in neural activity [40, 41]. Based on the in vitro study on mice in the neocortex, impairment in glutamate release, which is either due to the decreased function of glutamate receptors or loss of glutamate receptors causes the generalized absence and tonic-clonic epileptic seizures [42]. A study on the neocortex of mice [43] has shown that the paradoxical seizure exacerbation effect of the antiepileptic medication can be explained by the unintended suppression of inhibitory interneurons following the NMDA receptors blockade. Additionally, in an in vivo study on a mice model of tuberous sclerosis complex (TSC), the mutation in either TSC1 or TSC2 can disturb the function of NMDA receptors in the excitatory interneurons, which in turn can cause a shift from normal activity to ictal activity [15]. However, the mechanisms underlying the transition from interictal to ictal activities associated with glutamatergic receptors on EI remained unclear. Here, we take c_py-ei_ as the growing bifurcation parameter to investigate the dynamics of a model to explore the epileptiform activity induced by the glutamatergic receptor function of EI. According to Fig. 3, by gradually decreasing the EI excitability in the cortex, we can observe rich epileptiform transition dynamics.

**Fig. 3.**
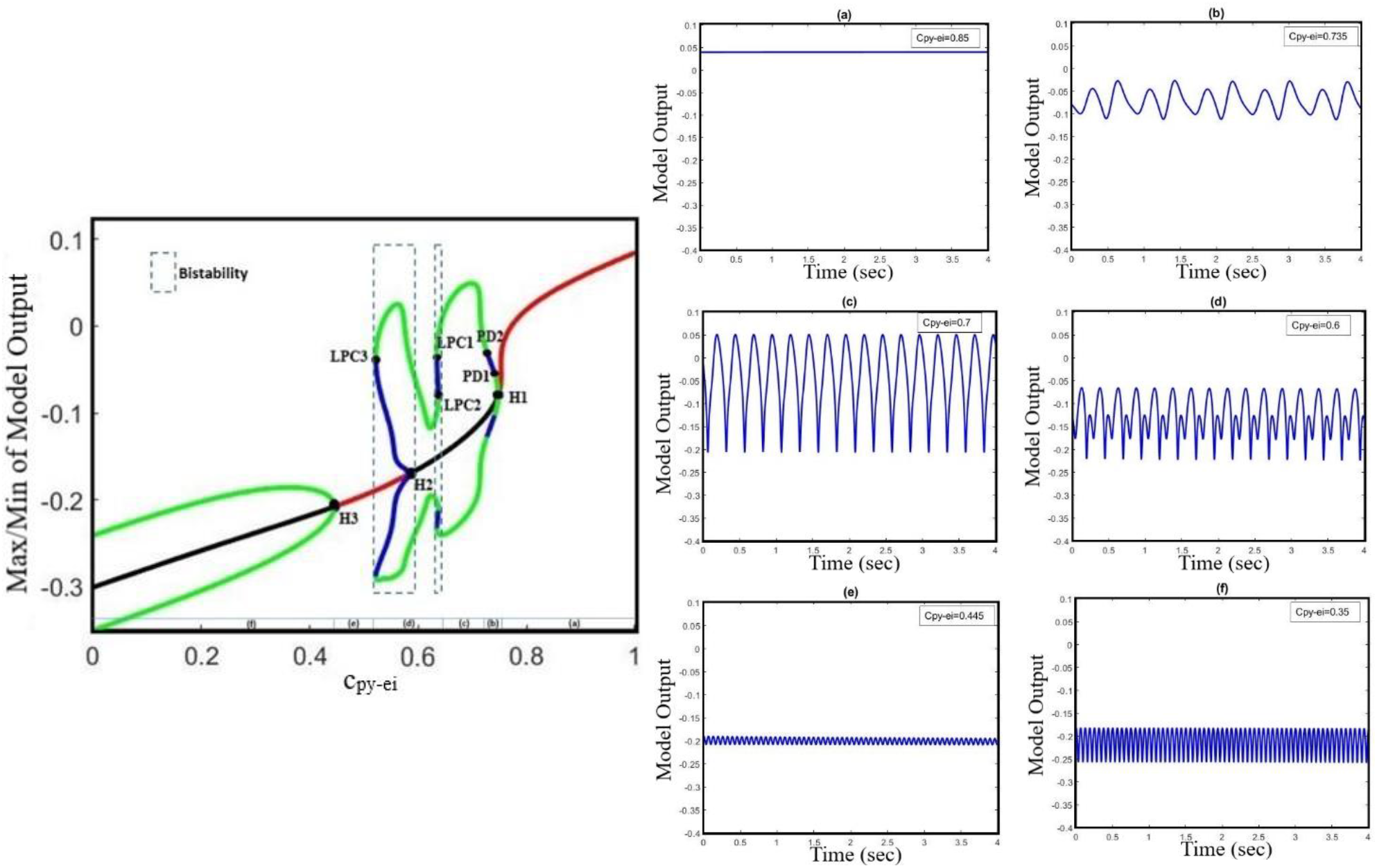
Bifurcation diagram and corresponding time series of the model output for different values of c_py-ei_. The bifurcation diagram of the model is calculated and plotted for the c_py-ei_ as the bifurcation parameter with c_i1-ei_ =0.3 and c_tc-ei_ = 4.5. We also set a_tc_=0 and a_py_=0. In the bifurcation plot, blue and green cycles, represent the unstable and stable limit cycles, respectively, and red and black lines, represent the stable and unstable points, respectively. It can be found from the bifurcation diagram that as the parameter c_py-ei_ decreases, (a) normal background firing (interictal), (b) pre-ictal spikes, (c) clonic discharges, (d) SWD, (e) slow rhythmic activity and tonic discharges can appear. LPC1, LPC2 and LPC3 are fold limit cycle bifurcation, H1 and H3 are supercritical Hopf bifurcation, H2 is subcritical Hopf, and PD1 and PD2 are period-doubling bifurcations.

According to the bifurcation diagram in Fig. 3, from right to left, we can observe the normal background activity, preictal spikes, clonic seizures, absence seizures, slow rhythmic activity and tonic seizures, respectively, combined with corresponding time series, as the parameter c_py-ei_ decreases. Hence, suppression of glutamatergic receptors on EI can lead to the transition from interictal to preictal signals, and also transition between epileptic seizures. With decreasing the synaptic strength c_py-ei_, the system encounters normal background activity (Fig. 3a) for the normal value of c_py-ei_. As the c_py-ei_ becomes smaller, a supercritical Hopf bifurcation point (H_1_) happens and the stable fixed points become unstable at c_py-ei_ ~0.74Upon c_py-ei_ is approaching to ~0.73, a period-doubling bifurcation (PD1) takes place and shapes the two amplitude preictal spikes pattern (Fig. 3b). Further decrease of c_py-ei_ parameter to ~ 0.72, another period-doubling bifurcation (PD2) happens and the starts to form clonic seizure patterns (Fig. 3c). In addition, with a further decreasing the c_py-ei_ parameter to ~ 0.678, a bistable region including two stable limit cycles takes place between the first- and second-fold limit cycle bifurcations (LPC1and LPC2), and shapes the SWD patterns (Fig. 3d). As the c_py-ei_ parameter decreases to ~ 0.591, the first subcritical Hopf bifurcation (H_2_) occurs and the stable points appear again. The second bistable region consists of stable focus points and a stable limit cycle happens between the subcritical Hopf bifurcation (H_2_) and the third fold limit cycle (LPC3) until c_py-ei_ is approaching to ~0.532. Upon decreasing the c_py-ei_ parameter, slow rhythmic activity (Fig. 3e) happens, and then at c_py-ei_ ~0.451 the second supercritical Hopf bifurcation (H_3_) arises from the steady-state and shapes the tonic seizure patterns (Fig. 3f).

Altogether, this model showed that impairment of GABAergic or glutamatergic receptors in the EI population causes the transition from normal background activity to seizures (preictal signal) and the transition between absence, tonic and clonic seizures. This model shows the possible existence of supercritical Hopf bifurcation with growing amplitude oscillations when the transition from interictal to ictal state occurs. In order to have a deeper understanding of the interictal to ictal transition dynamics, we investigate the bifurcation of the cortical and sensory input frequencies in our model.

#### 3.1.3. Hybrid cooperation of GABAergic and glutamatergic receptors of EI in epileptic transition dynamics

The genesis of generalized seizures requires interdependencies in different thalamocortical connections. Studies on genetic rat models have shown that the increasing excitatory coupling strength from thalamus to cortex may facilitate the maintenance and propagation of absence seizures [44,45]. However, based on the paradoxical behavior of glutamatergic and GABAergic receptors discussed in sections 3.1.1 and 3.1.2, here, we explore the effect of decreased glutamatergic synaptic strength from the thalamus to neocortex on seizure propagation using the extended model. In this section, we investigate the effect of a dual collaboration of intra-EI GABAergic and glutamatergic synaptic strength in the cortex on the transition between different kinds of activity, either from interictal to ictal signal or from absence seizure to tonic and clonic seizures. Moreover, we explore the influence of TC to EI pathway dominated by glutamatergic receptors on the initiation and propagation of epileptic activities based on the altered ratio of dual cooperation of intra-EI GABAergic and glutamatergic receptors in the cortex. According to Fig. 4, the pattern evolutions for different values of c_tc-ei_ are obtained to investigate the state transition produced by this model and their corresponding dominant frequency distributions as the parameters of c_py-ei_ and c_i1-ei_ varies in region [0.1,0.9] × [0.2,0.9].

**Fig. 4.**
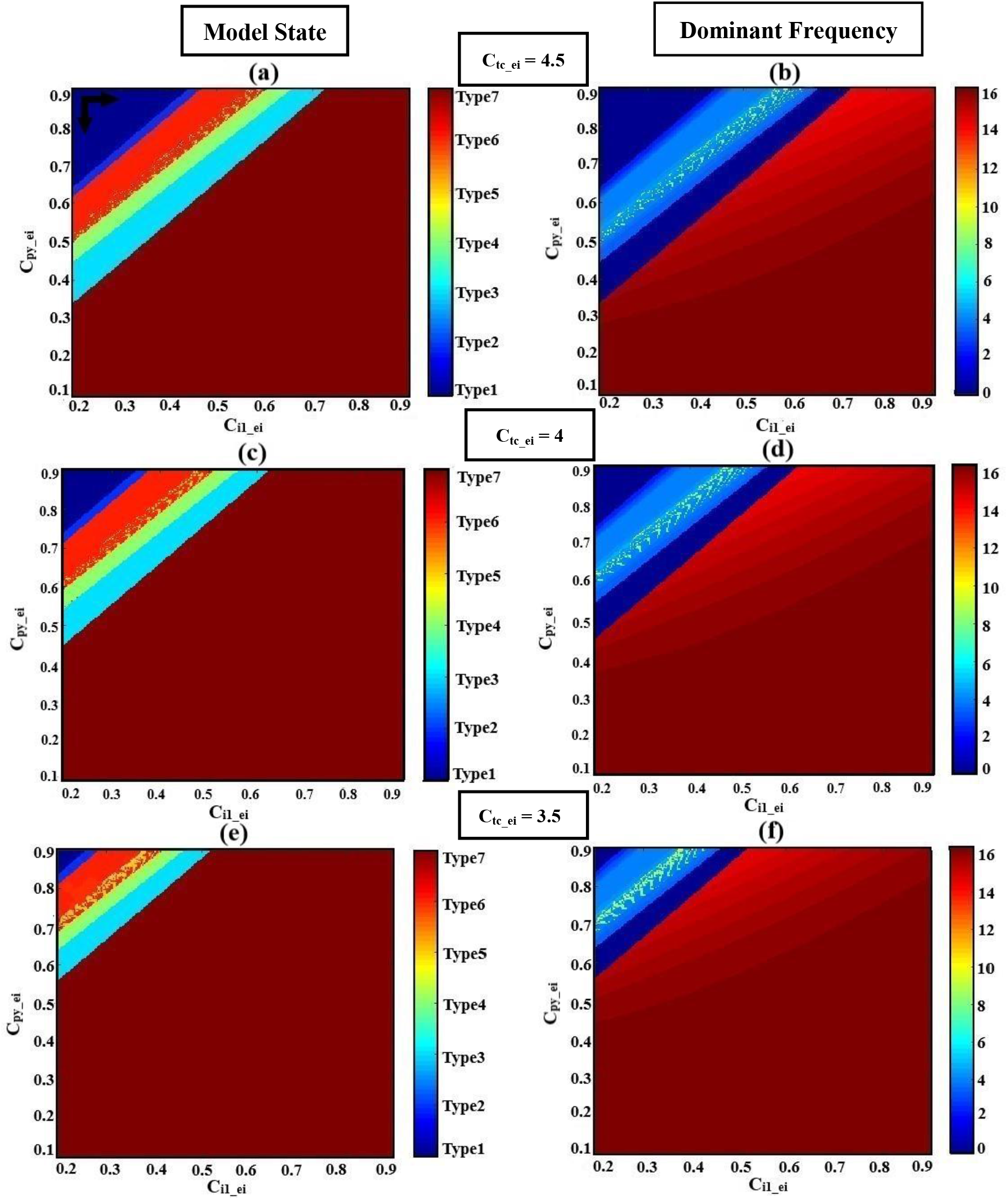
Hybrid modulation of the model. Patterns of evolutions of the model states are shown by changing the glutamatergic and GABAergic synaptic strength of EI (c_py-ei_, c_i1-ei_) in the cortex. Each row corresponds to a given TC to EI excitatory synaptic strength (c_tc-ei_). The left column shows the states (normal or epileptic activities) and the right column shows the dominant frequencies (DF) of the model output. We set the a_tc_=0 and a_py_=0. The various activities **(a), (c)** and **(e)** and their corresponding dominant frequencies **(b), (d)** and **(f)** with decreasing the excitatory synaptic strength c_tc-ei_ from top to bottom, 4.5, 4 and 3.5, respectively, are shown. Model produces different firing sates with variation of c_py-ei_, c_i1-ei_. These states are normal background firing (Type1; DF = 0 Hz), preictal activity (Type2; DF < 3.5 Hz), slow rhythmic activity (Type3; DF = 0 Hz), typical absence seizure (Type4; 2 Hz < DF < 4 Hz), atypical absence seizure (Type5; DF > 4 Hz and DF < 2 Hz), clonic seizure (Type6; DF ≤ 7 Hz) and tonic seizure (Type7; DF ≥ 14 Hz). The black horizontal and vertical arrows represent the transition routes from the normal state to the pathological states.

For higher values of synaptic strength like c_tc-ei_ =4.5 (Fig. 4(a) and (b)), this model demonstrates seven different types of activities, when c_i1-ei_ varies from 0.2 to 0.9. We denoted these seven activities as follows. type1, normal background activity (DF = 0 Hz in Fig. 4(b)); type2, preictal activity (DF < 3.5 Hz in Fig. 4(b)); type6, clonic seizure (DF ≤ 7 Hz in Fig. 4(b)); type5, atypical absence seizure (DF > 4 Hz and DF < 2 Hz in Fig. 4(b)); type4, typical absence seizure (2 Hz < DF < 4 Hz in Fig. 4(b)); type3, slow rhythmic activity (DF = 0 Hz in Fig. 4(b)); type7, tonic seizure (DF ≥ 14 Hz in Fig. 4(b)). Upon decreasing the synaptic strength c_tc-ei_ from 4.5 to 4 (Fig. 4(c) and (d)), one can see that the surface of the yellow region (atypical absence seizure; Type5) and red region (tonic seizure; Type7) increase with the reduction level of excitation of intra-EI. Accordingly, the regions correspond to normal background activity (Type1), preictal activity (Type2), clonic seizure (Type6), typical absence seizure (Type4) and slow rhythmic activity (Type3) decreases. Biologically, decreasing the excitatory coupling strength from TC to EI interrupts the normal activity of the thalamic relay nucleus in the thalamocortical network. This interruption facilitates the transition from interictal to ictal, *i.e*., with the increasing of c_py-ei_ and c_i1-ei_ from their normal state, the transition from normal background activity to ictal activity and also the transition between clonic, absence and tonic seizures can occur more easily. Upon further decreasing of the c_tc-ei_ to 3.5 (Fig. 4(e) and (f)), the yellow and red regions correspond to atypical absence seizure (Type5) and tonic seizure (Type7) extend, respectively, However, the whole other regions decrease, which shows that the transition from normal background activity to epileptic seizure activities in this model can be influenced by the intra-EI GABAergic and glutamatergic receptors in the cortex under the impact of the thalamus to the cortex synaptic strength.

In summary, the intra-EI GABAergic and glutamatergic receptors in the cortex can elicit seven different kinds of activity in this model under the effect of excitatory synaptic strength from TC to EI. The lower excitatory synaptic strength of c_tc-ei_, in the model, causes a faster transition from interictal to ictal state. This implies the important role of the thalamus to cortex synaptic strength in the initiation and maintenance of generalized epileptic seizures.

#### 3.1.4. Transition dynamics produced by glutamatergic receptor-mediated excitation in RE

Increasing the GABAergic inhibition is predominantly believed to suppress epileptic seizures, however, studies [46, 47, 48] have reported that enhanced GABAergic inhibition in the brain promotes seizure as well. The reticular nucleus (RE) is mainly composed of inhibitory interneurons that their epileptogenic role in seizure is investigated by Liu and Wang in their thalamocortical model [22]. Liu and Wang did not consider the EI population in their model when they investigated the role of TC to RE synaptic strength in epileptic activities. Therefore, here, we investigate how does the strength of the excitatory synapses of TC affect the function of the inhibitory reticular nucleus population in seizure generation in presence of EI. By increasing the strength of the excitatory synapses from TC to RE, we increase the inhibitory behavior of RE in the proposed thalamocortical model. In Fig. 5 we show that the proposed model demonstrates a rich dynamic and that the transition from interictal to ictal occurs when we increase the ctc-re. In Fig. 5, from left to right, first, we observe the normal background activity. After increasing the parameter c_tc-re_ to ~ 9.4, the transition from steady-state to the limit cycle of preictal (supercritical Hopf bifurcation H1) happens. Further increasing the c_tc-re_ to ~ 9.58 and 9.63 the first and second period-doubling bifurcations (PD1, PD2) take place, respectively. After PD2, the limit cycles correspond to clonic seizure take place. With increasing the c_tc-re_ to ~ 10.1 the first fold limit cycle (LPC1) and then at c_tc-re_ ≈ 10.3 the second fold limit cycles (LPC2). the model generates the first bistable region between LPC1 and LPC2, which shape the limit cycle corresponding to absence seizure activity. The second bistable region occurs between the subcritical Hopf bifurcation H2 at ~ 10.6 and the third fold limit cycle LPC3 at ~ 11.9. Then, at c_tc-re_ ≈ 11.8 the transition from limit cycles of absence seizures and steady-state occurs. Upon increasing the c_tc-re_ to ~12.1, the transition from the stable equilibrium to the limit cycle of tonic seizure happens. By increasing the parameter c_tc-re_ to 9.5, we can observe the first supercritical Hopf bifurcation (H_1_) and the transition from interictal to preictal state. Then by further increasing the c_tc-re_ and inducing more impairment in the TC to RE pathway, we can observe limit cycles correspond to clonic and absence seizures, respectively.

**Fig. 5.**
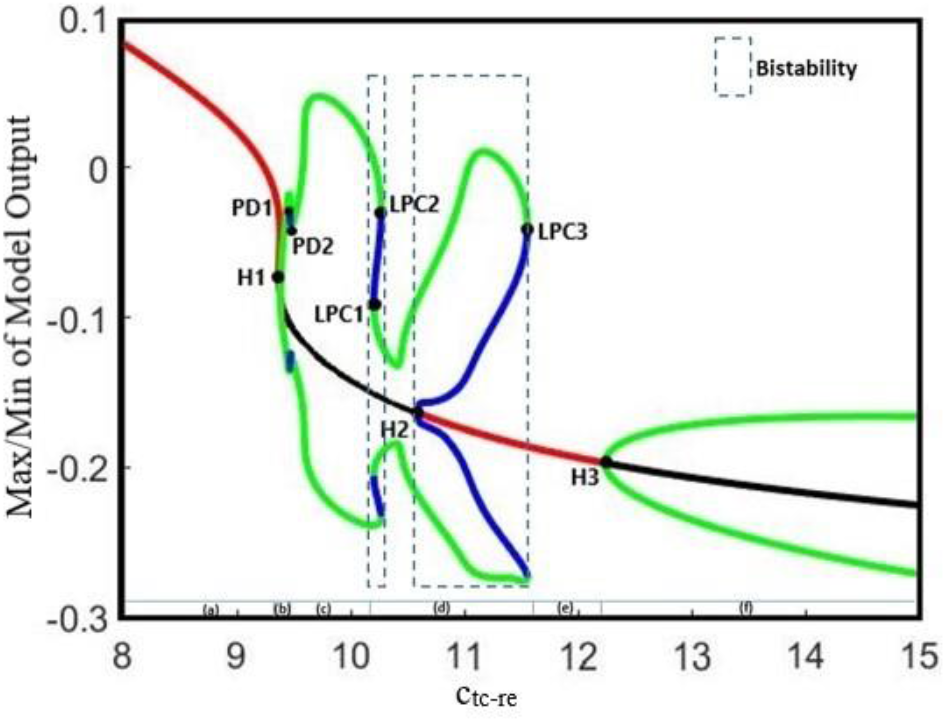
The bifurcation diagram for the bifurcation parameter c_tc-re_. Other parameters of the model are c_py-ei_ =0.75, c_i1-ei_ =0.33 and c_tc-ei_ = 4.2. We also set a_tc_=0 and a_py_=0. In the bifurcation diagram, blue and green cycles represent the unstable and stable limit cycles, respectively. The red and black lines represent the stable and unstable points, respectively. LPC1, LPC2 and LPC3 are fold limit cycle bifurcation, H1 and H3 are supercritical Hopf bifurcation, H2 is subcritical Hopf, and PD1 and PD2 are period doubling bifurcations.

### 3.2. Frequency analysis of the different initial states of the model

According to clinical observations, sensory [49] and cortical [50] stimulations can provoke epileptic seizures. Therefore, in this section, we examine the effect of sensory and cortical inputs in the interictal to ictal transition using the extended thalamocortical model. Studies on thalamocortical models [51, 52], have shown that changes in the frequencies of cortical and sensory periodic inputs can cause a transition from chaotic to periodic activities in the output of the model. Therefore, we will first examine the effect of frequencies of the cortical and sensory inputs on the transition behavior of our model, and even more importantly, we will discuss the effect of the initial state of the model on this transition.

Fig. 6, shows various bifurcation diagrams for different initial states of the model, such as, normal background activity (interictal), preictal activity, clonic, absence and tonic seizures. To change the initial state of the model, we used different values for parameter c_py_ei_. These values were selected using the results of the bifurcation analysis shown in Fig. 3, in which c_py_ei_ is the bifurcation parameter. In Fig. 6(a) and (b), we set the c_py_ei_=0.76, which is in the range corresponding to interictal activities. In Fig. 6(c) and (d) by decreasing the parameter c_py_ei_ to 0.748, the initial state of the model corresponds to preictal activities. In the Fig. 6(e) and (f), we change the initial state of the model to a clonic seizure state by decreasing the value of c_py_ei_ to 0.72. As it is shown in Fig. 6(g) and (h), and Fig. 6(i) and (j), by further decreasing the c_py_ei_ to 0.58 and then to 0.4, we set the model in absence seizure and then in tonic seizure states, respectively.

**Fig. 6.**
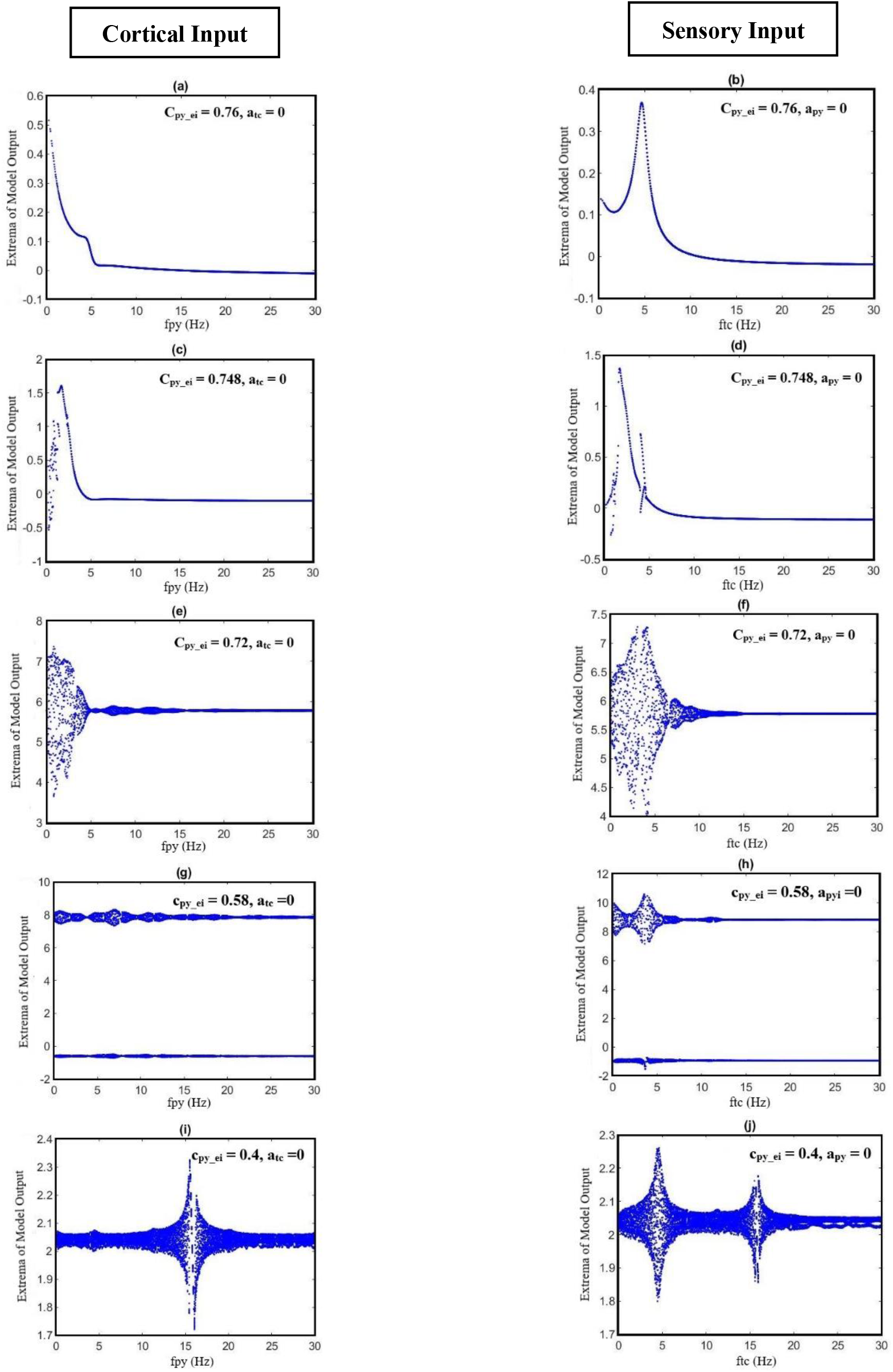
The bifurcation diagrams as each parameter of fpy and ftc changes. The various bifurcation diagrams of the model output when the frequencies of the cortical input (fpy) and sensory input (ftc) change. Other parameters have the numerical values of c_i1-ei_ =0.3 and c_tc-ei_. We set the a_tc_ = 0 as we change the fpy in diagrams in the left colums. In diagrams in the right colums the ftc changes and the apy is set equal to zero.

On the other hand, for each state of the model, we changed the frequencies of cortical and sensory periodic inputs separately with a frequency step increment of 0.01 Hz. For the interictal initial state (Fig. 6(a) and (b)), the model shows a linear resonant behavior with a resonant frequency of fpy ≈ 0.2 and ftc ≈ 4.7 as the cortical and sensory input frequencies increase. The model, in this case, behaves like a bandpass filter for the sensory input and a low pass filter for the cortical input. For c_py_ei_ = 0.748, when the frequency of cortical input is increased, a chaotic behavior is resulted. In Fig. 6 (c), further increase in the cortical input frequency (fpy), we notice a jump near the frequency of fpy ≈ 1.7. Moreover, by increasing the frequency of the sensory input, two jumps take place near the frequencies of ftc ≈ 1.8 and ftc ≈ 4 (Fig. 6(d)). Then, upon increasing the fpy and ftc, model returns to its normal periodic behavior. This observation indicates that, when initial state of the model changes from normal background activity to preictal activity, the behavior of the model changes from a linear resonator to a nonlinear resonator. In Fig. 6 (e) and (f), where the model is in the clonic seizure state, by increasing the frequencies of cortical and sensory inputs, we observe that the model first displays a noticeable chaotic behavior around 3 Hz, which is the main frequency of the clonic seizure, and then returns to its expected periodic behavior. In Fig. 6 (g) and (h), when model is in the absence seizure state, increasing the cortical and sensory input frequencies yield two chaotic behaviors with two different amplitude ranges; the reason for this is the coexistence of the two different SWD patterns with different amplitudes. This chaotic behavior can be seen around the main frequency of absence seizure (fpy ε [2, 4]). As we can see in Fig. 6 (i) and (j), when the state of the model changes to tonic seizure activities, we observe chaotic behavior around the main frequency of tonic seizure (fpy ≈ 16). As it is demonstrated in Fig. 6 (f), (h) and (j), when the model is in seizure activity states, by increasing the frequency of sensory input, two peaks in the frequency response of the model is observed. These two peaks can be seen more clearly in tonic seizures (Fig. 6 (j)), and correspond to the two jumps already observed in Fig 6 (d).

Fig. 7 demonstrates the effect of the initial state of the model on the transition between small amplitude oscillation and large amplitude seizure activities, all in the time domain. We set the c_py_ei_ equal to 0.76 in order to adjust the model in its normal state and then we change the model state from interictal (normal activity) to preictal activity by decreasing the c_py_ei_ parameter to 0.748. In Fig. 7(a), the model receives a cortical input with a fixed frequency, fpy=1 Hz, in the absence of sensory input. However, in Fig. 7(b), we only consider the effect of the sensory input on the output of the model and we set its frequency ftc = 1 Hz. In Fig. 7(a) and (b), it is shown that based on the bifurcation diagram of Fig. 3, by changing the c_py_ei_ from 0.76 to 0.748, the state transition from normal activity to typical absence seizure activity occurs, and then it returns to the normal activity as c_py_ei_ returns to its initial value.

**Fig. 7.**
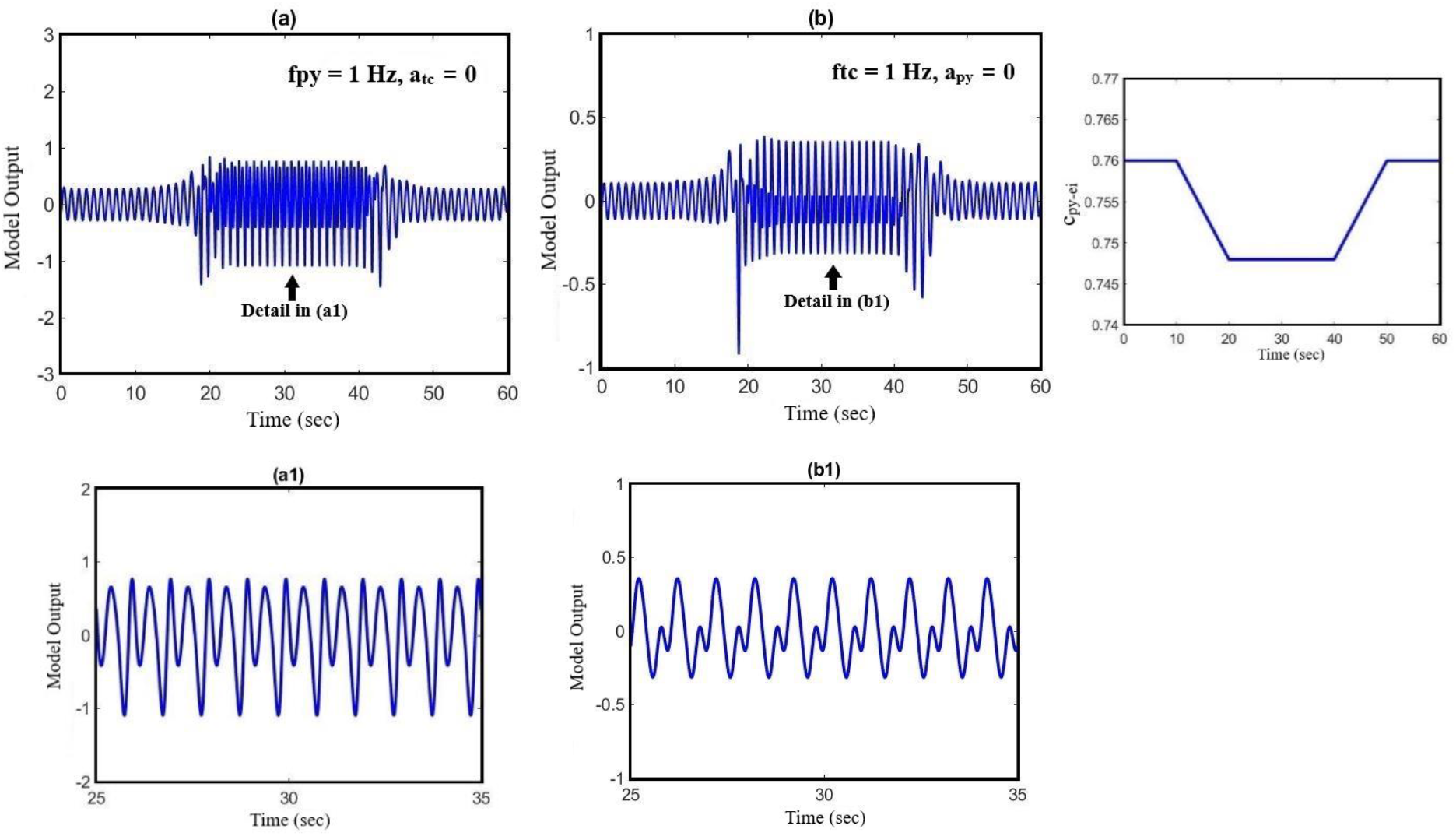
The effect of state transition of the model on its output when the parameter of c_py_ei_ changes. The effect of initial state of the model on the transition between periodic signal and seizure when the cortical and sensory input frequencies are constant. In (a) we set the parameters of fpy = 1 Hz and atc = 0, and In (b) we set ftc=1 HZ and a_py_=0. The pattern of glutamatergic synaptic strength, c_py_ei_, is presented on the right-hand side diagram.

## 4. Discussion

Traditionally, mechanisms underlying seizures have been considered to be due to increased excitation, decreased inhibition or even both of them resulting in hyper-excitability and seizure generation. However, glutamatergic and GABAergic neurotransmitters can exert paradoxical effects and cause seizures. Antiepileptic drugs do not necessarily work by decreasing excitation and increasing inhibition of neural activities. Studies [53–57] have shown that some seizures occur when inhibition is enhanced in the brain. Therefore, it might be oversimplifying if one considers the role of GABAergic inhibition in the brain as the only antiepileptics. In fact, understanding the seemingly contradictory role of the excitatory and inhibitory neurons in the brain can lead to new therapies for epileptic seizures. Therefore, in the present study, we investigated the opposite effect of GABAergic and glutamatergic receptors in seizure onset through the dynamical analysis of an extended model. In this direction, we proposed an extended neural mass model, which considers the role of the spiny stellate cell population connectivity with other cortical neural populations in different states of seizure generation. The original thalamocortical model, which was developed by Liu and Wang in 2017 [22], was modified and extended and then used to simulate the preictal activity that has a prevailing role in epileptic seizure prediction. Using this model, we investigated the interactions between excitatory interneurons and the glutamatergic pyramidal and inhibitory interneurons in the cortex, and how they lead to epileptic seizures. This model enabled us to generate preictal activities before the clonic seizure. To make this point clearer, we calculated the bifurcation diagram of the output of the model for **c_i1-py_**, as the bifurcation parameter, in two cases: 1) without EI population in the model, and 2) at the presence of EI. As one can see in both Fig. 8 (a) and (b) by increasing the inhibitory synaptic strength between the PY and I1, first clonic seizure and then, a tonic seizure occurs. In Figure (a), when the EI population is not considered in the model, there exist no preictal activities (please notice the high amplitude clonic activity after Hopf bifurcation H1). However, in Figure (b), when the role of excitatory interneurons in the model is considered, the preictal activity emerges in the bifurcation diagram right after Hopf bifurcation H1 (please notice the low amplitude preictal activity after H1). Therefore, one can conclude that although the order and list of bifurcations that occur when the inhibition between fast interneurons and pyramidal neurons increases, are the same for both cases (extended model and the original model developed by Liu and Wang [22]), but the emergence of the preictal activities in the model depends on the excitatory interneurons. Without these neurons (and certainly the feedback loop created by them) the model does not show any preictal activities and the seizure starts abruptly.

**Fig. 8.**
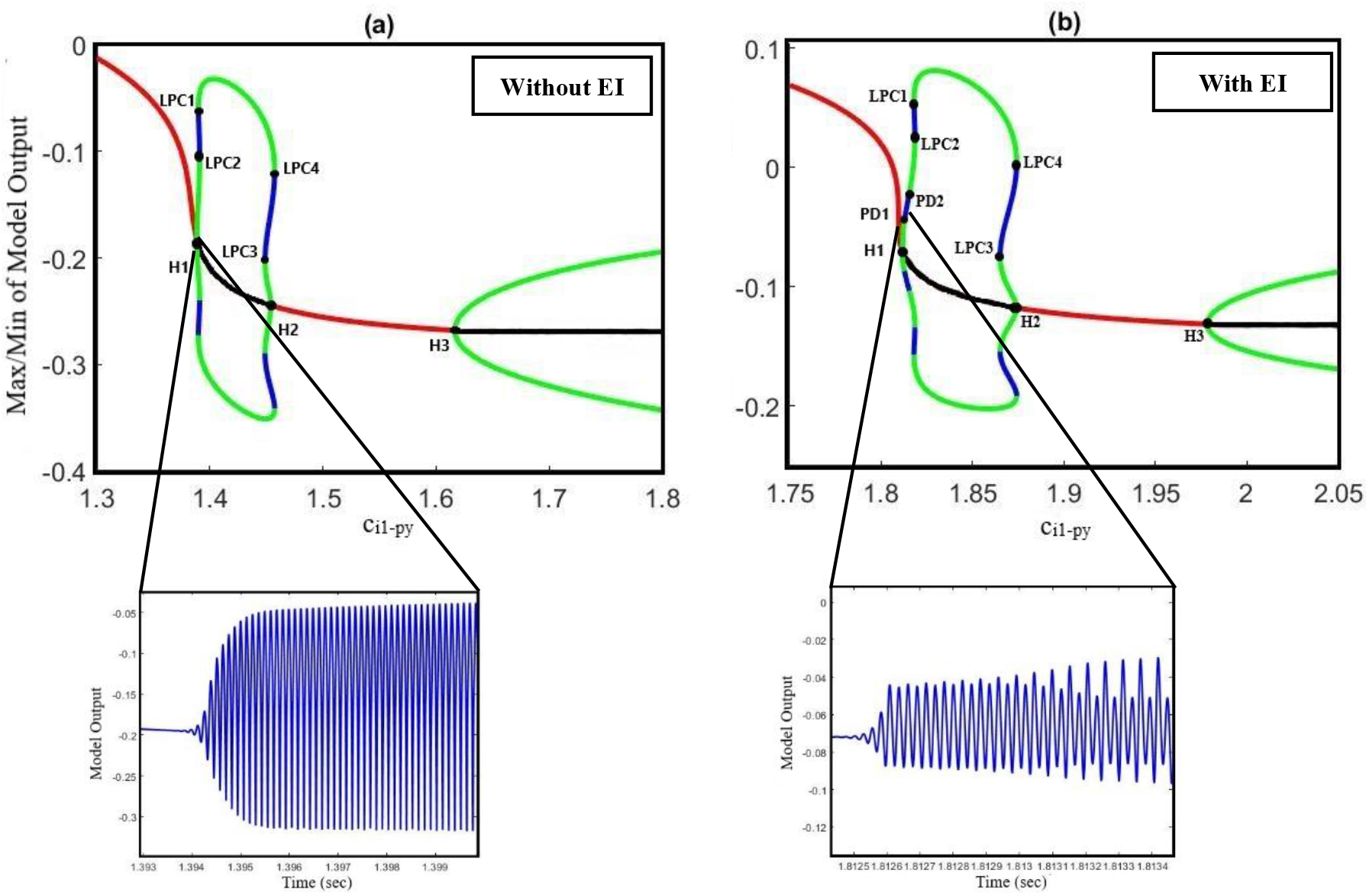
(a): Bifurcation diagram of the output of the model when the excitatory interneurons are not considered in the model (parameters are set based on the caption of Fig.1). Bifurcation parameter is the inhibition strength between I1 (fast interneurons) and PY (pyramidal) population (c_i1_py_). (b): Bifurcation diagram of the output of the model when the excitatory interneurons are considered in the model (c_py_ei_= 0. 8, c_i1-ei_=0.3, c_tc-ei_=4.5). Bifurcation parameter is the inhibition strength between I1 (fast interneurons) and PY (pyramidal) population (c_i1_py_).

Our results show the richness of the dynamics that the proposed model can generate, including normal background activity, preictal spikes, slow rhythmic activity, clonic seizure, typical absence seizure and tonic seizure as we decreased and increased the glutamatergic and GABAergic synaptic strengths of the EI, respectively. Furthermore, bifurcation diagrams of the model were obtained by varying the coupling strength of GABAergic and glutamatergic receptors in EI. The diagrams in Fig.2, Fig.3 and Fig.5 show that the interictal to ictal transition occurs when we increased the glutamatergic excitation and GABAergic inhibition in the cortex. Based on our bifurcation diagrams, pathological transitions are consequences of supercritical and subcritical Hopf bifurcations as well as the fold limit cycle bifurcation. The onset of preictal discharges is characterized by a supercritical Hopf bifurcation. This; results in a preictal activity, which is characterized by growths in amplitude and frequency of neural activities. As mentioned before, Liu and Wang [22] developed a thalamocortical model, which is originally inspired by Taylor et al. [21] and Fan and colleagues [17] to study the epileptogenic role of the synaptic strength between TC and ER in the thalamus. Using that model, they did not consider the spiny stellate cells population and its role in epileptic seizure generation. One of the most important features lacking in their results was the preictal state of the model. In fact, the model was not able to generate the limit cycles of preictal state in their bifurcation analysis when they investigated the excitatory pathway from TC to RE. As a result, in their bifurcation diagrams there was an abrupt appearance of ictal limit cycles after the normal background activities. However, in Fig.5 we evaluated the extended thalamocortical model in a more general framework and we showed that this model is capable of generating the preictal state when the synaptic strength from TC to RE is changed. We showed that considering the role of excitatory interneurons was crucial for the emergence of the preictal state. Using this model, we demonstrated that ictal discharges do not appear abruptly after a period of interictal activities. As it was shown in the bifurcation diagrams (Fig. 2, 3 and 5), we can simulate the gradual increase in frequency and amplitude from normal activities to ictal activities. Consequently, the model demonstrates the key role of the spiny stellate cells of cortex in interictal to ictal transition dynamics.

In order to have a deeper insight into the dynamics of interictal to ictal transition, we also explored the effect of cortical and sensory periodic (sinusoidal) inputs on the output of the extended and modified model. Our simulation results reveal that the transition from normal activity to seizure activity occurs when a perturbation in the dynamic structure of the model relocates the model state from interictal to preictal state. Our results show that during the preictal state, the extended model is more susceptible to sensory and cortical inputs, which in turn, causes typical absence seizures. Moreover, the frequency response of the model (Fig. 6) makes it clear that model responses to the cortical and sensory input are dependent on the initial state of the model. According to the bifurcation analysis depicted in Fig. 6, when the model is in its normal state, it behaves as a linear resonator. Therefore, when we stimulate the model with sensory and cortical stimuli, the model output doesn’t show a chaotic behaviour and comes back to its normal behavior as the input cortical and sensory frequencies are increased. On the other hand, when any impairments occur in the glutamatergic receptors of the excitatory interneurons, these cause the model to enter the preictal state. Then, the same sensory and cortical stimulations can cause the model to act as a nonlinear resonator and we can observe chaotic behaviors and jump phenomenon. This nonlinear behavior increases when the initial state of the model changes to the ictal state. In general, the cortical stimulations evoke more peak to peak amplitude of cortical output in this model, in comparison to sensory stimulations, when it is in clonic and tonic seizures states. On the contrary, the peak to peak amplitude of the model output when it is in seizure activity state is more intense when it receives sensory stimulations. This demonstrates that absence seizures are more sensitive to sensory stimuli than cortical stimulations. Also, we investigated the effect of the initial state of the model in absence seizure generation. As it can be seen in Fig.7, when the extended model is in its normal state, we observe the low amplitude activities, which correspond to normal background activity. However, by changing the model state from normal to preictal state, we can observe that the absence seizures take place. According to our observation, we suggest that alteration in the initial states of the brain can be considered as one of the principal causes of the absence and photosensitive seizures.

Finally, we examined the role of the thalamus in epileptic seizure activities. We investigated the role of cooperation of glutamatergic and GABAergic receptors of EI under the effect of the thalamic relay nucleus (TC) in seizure generation and propagation. As it can be observed in Fig.4, the results have shown the dependence of cortical function on the thalamic one in initiation, propagation and termination of epileptic seizures Recent studies [58, 59] did not consider the thalamus and its synaptic connectivity with other neuronal populations such as excitatory interneurons in the cortical models they worked with. However, based on the bifurcation analysis obtained from hybrid modulation of intra-EI excitatory and inhibitory synaptic strength in the extended neural mass model, the pathway from thalamic relay nucleus to the excitatory interneurons facilitates the transitions between different types of epileptic activities, either from interictal to ictal or between clonic, absence and tonic seizures. We believe these results provide useful insights for understanding more thoroughly the function of EI population and dynamics caused by their connections to other populations in the thalamocortical circuitry and the transitions between inter-ictal to preictal and ictal states. Therefore, the extended model is more suitable to be used in epileptic seizure prediction and abatement.

